# Acute Metabolic Switch by High-Dose Dehydroascorbic Acid: Rapid TCA Cycle Depletion and Early Signature of Methionine Trap within 1 Hour in KRAS-Mutant Cells

**DOI:** 10.64898/2026.10.01.755313

**Authors:** Nozomu Hatakeyama, Taiki Miyazawa, Masako Toda, Teruo Miyazawa

**Author notes:** **Corresponding author:** Nozomu Hatakeyama.

## Abstract

Dehydroascorbic acid (DHA), the oxidized form of vitamin C, exhibits specific toxicity against KRAS-mutant cancer cells; however, the early metabolic reprogramming mechanisms leading to cell death remain incompletely understood. In this study, we reveal the non-linear, dose-dependent response of the intracellular metabolome during the ultra-early phase of 1-hour DHA exposure. Under 1 mM DHA exposure, an early significant accumulation of the primary catabolite erythrulose (*p* = 0.030) and moderate consumption of glutamate (*p* = 0.0086) were observed, while core TCA cycle homeostasis and cell viability were preserved. In contrast, under 10 mM DHA exposure, an acute and profound alteration of the metabolic profile was observed. Along with the significant accumulation of glycerate, indicative of a glycolytic blockade, and the terminal peroxidation product oxalate, core TCA cycle intermediates such as α-ketoglutarate and malate showed rapid depletion (Log_2_FC < –2.0, *p* < 10^−7^). Furthermore, coupled with this acute energy depletion, a statistically significant accumulation of free methionine (Log_2_FC = 1.14, *p* = 0.011), which was below the limit of detection in the control, was identified alongside profound depletions of glutamate (*p* < 0.001) and cysteine (*p* = 0.0027). These results strongly suggest that the initial response to high-dose DHA is not merely a linear accumulation of oxidative stress, but rather an “abrupt metabolic switch” characterized by rapid TCA cycle dysfunction followed by the early signature of a methionine trap.

## 1. Introduction

KRAS-mutant cancer cells overexpress glucose transporters (e.g., GLUT1) and possess the property of specifically and massively taking up dehydroascorbic acid (DHA), the oxidized form of vitamin C (Agus et al., 1999; Yun et al., 2009). During the reduction of internalized DHA back to ascorbic acid (Asc) by antioxidants such as glutathione (GSH), intense oxidative stress (reactive oxygen species, ROS, generation) and the depletion of the NADPH/GSH pool occur, ultimately inducing cell death (Yun et al., 2015). Previous studies have shown that at DHA exposures of 1 mM or lower, cells can dose-dependently and linearly reduce and accumulate DHA as ascorbic acid using antioxidant networks (Abe et al., 2022). However, much remains unknown regarding where the first decisive breakdown occurs within major intracellular metabolic networks (glycolysis, TCA cycle, and amino acid metabolism) during the “ultra-early phase” (tens of minutes to 1 h) when an ultra-high dose (10 mM) of DHA, which transcends this reduction capacity limit (threshold), influxes into the cells. This study conducted a comprehensive metabolomic analysis of KRAS-mutant cells 1 h post-DHA exposure. By comparing a low dose (1 mM) and a high dose (10 mM), we clarify the boundary between the homeostatic maintenance phase in response to DHA influx and the acute metabolic collapse phase following threshold breakthrough, thereby reporting the full picture of the “early metabolic switch” that serves as the starting point of the antitumor effect.

## 2. Results

### 2.1. Initial Response to 1 mM DHA Exposure and Non-linear Glycolytic Blockade at 10 mM Exposure

The human gastric adenocarcinoma cells (AGS cells) have been reported to harbor an activating KRAS G12D mutation (Kneissl et al., 2012). We therefore selected AGS cells as a cancer model with a previously reported KRAS mutation to characterize the early metabolic response to DHA exposure. To evaluate the dose-dependent threshold response of DHA uptake and initial metabolism of AGS cells, we compared the intracellular metabolomes 1 h after 1 mM (*n* = 11, across 2 batches) and 10 mM (*n* = 18, across 3 batches) DHA exposure (**Figure 1**). Erythrulose, an early cleavage product of DHA, already showed a statistically significant ~3.1-fold accumulation (Log_2_FC = 1.64, *p* = 0.030) in the 1 mM exposure group compared to the control group, and an even more pronounced 9.3-fold accumulation (Log_2_FC = 3.22, *p* = 8.32 × 10^−9^) in the 10 mM group (**Figure 2, top row**). This indicates that intracellular influx and initial breakdown of DHA proceed even at a low dose of 1 mM without crashing the cell system.

**Figure 1.**
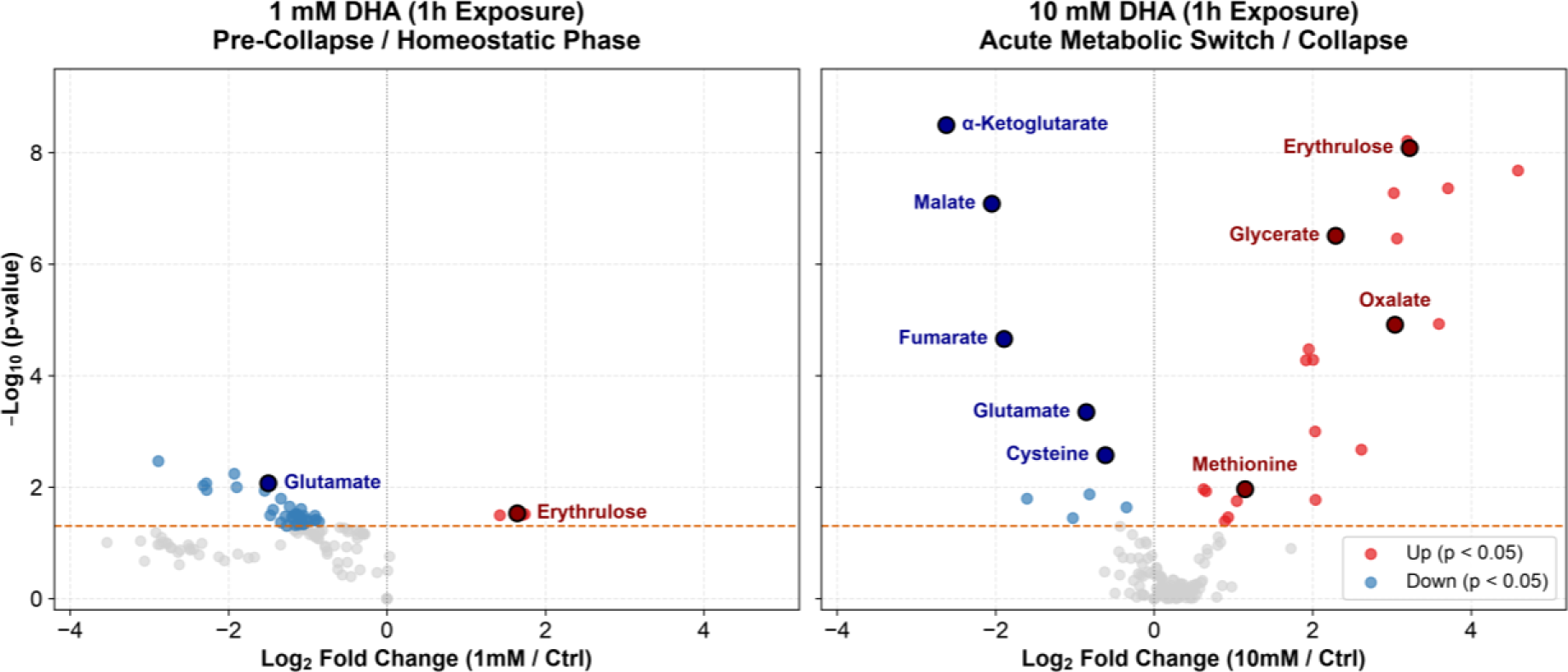
Non-linear metabolic responses of AGS cells to 1 mM and 10 mM DHA exposure (1 h). Volcano plots highlighting the global metabolomic changes in cells exposed to 1 mM (left, *n* = 11) and 10 mM (right, *n* = 18) DHA for 1 h, compared to untreated controls. The x-axis represents the Log_2_ fold change, and the y-axis represents the –Log_10_ (*p*-value). The horizontal dashed orange line indicates the significance threshold (*p* = 0.05). While 1 mM DHA induced mild accumulation of early catabolites such as erythrulose without perturbing core TCA homeostasis, 10 mM DHA triggered acute depletion of TCA cycle intermediates and specific amino acids (blue dots, e.g., α-ketoglutarate, malate, glutamate, cysteine) and massive accumulation of DHA breakdown products and specific amino acids (red dots, e.g., oxalate, glycerate, methionine).

**Figure 2.**
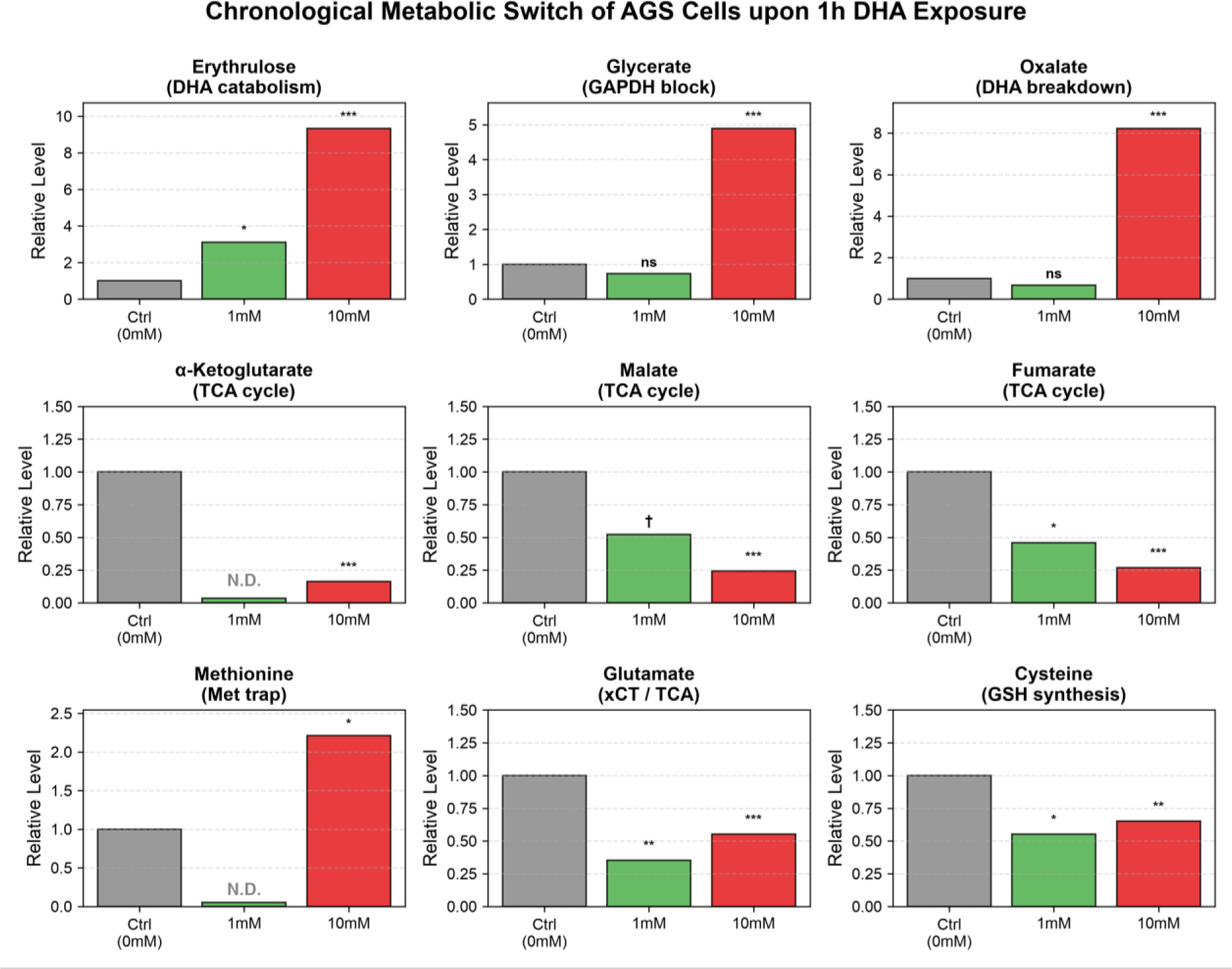
Chronological metabolic switch of AGS cells and early signature of the methionine trap upon DHA exposure. Relative levels of key metabolites arranged in chronological sequence of the metabolic cascade: DHA catabolism and glycolytic blockade (**top row**), TCA cycle arrest (**middle row**), and amino acid dynamics / methionine trap (**bottom row**). Data represent mean relative abundance normalized to the control group (0 mM = 1.0). While 1 mM DHA (*n* = 11, pooled from 2 independent experiments) induced mild accumulation of early catabolites (erythrulose) without disturbing core TCA cycle integrity, 10 mM DHA (*n* = 18, pooled from 3 independent experiments) triggered a cascading metabolic failure characterized by acute TCA depletion, massive accumulation of glycerate/oxalate, and the specific accumulation of methionine. Asterisks and symbols denote statistical significance versus control (*** *p* < 0.001, ** *p* < 0.01, * *p* < 0.05, †*p* < 0.10 [statistical trend], ns: not significant). N.D.: Not Detected.

Conversely, oxalate, a terminal product of non-enzymatic DHA peroxidation (Simpson & Ortwerth, 2000; Linster & Van Schaftingen, 2007), and glycerate, an indicator of GAPDH inactivation, exhibited a completely non-linear response profile. Both metabolites remained without significant alteration in the 1 mM group (Oxalate: FC = 0.68, *p* = 0.380; Glycerate: FC = 0.73, *p* = 0.403), but exhibited explosive, massive accumulations of 8.2-fold (Log_2_FC = 3.04, *p* = 1.22 × 10^−5^) and 4.9-fold (Log_2_FC = 2.29, *p* = 3.10 × 10^−7^), respectively, in the 10 mM group (**Figure 2, top row**). These non-linear dynamics strongly indicate that 10 mM DHA exposure breaches the intracellular antioxidant threshold, triggering acute glycolytic blockade and uncontrolled DHA auto-degradation.

### 2.2. Rapid Collapse of the TCA Cycle Under 10 mM DHA Exposure

To assess the downstream energetic consequences of this blockade, we analyzed mitochondrial TCA cycle intermediates. In the 1 mM exposure group, core TCA cycle intermediates maintained overall metabolic homeostasis without catastrophic collapse (α-ketoglutarate remained unchanged, while malate and fumarate showed mild, modest decreases: Malate: Log_2_FC = –0.94, †*p* = 0.068; Fumarate: Log_2_FC = –1.12, *p* = 0.047) (**Figure 1, Left; Figure 2, middle row**).

In sharp contrast, 10 mM DHA exposure caused a rapid, systemic collapse of the TCA cycle within 1 h: α-ketoglutarate plummeted to less than 1/6 of control levels (Log_2_FC = –2.62, *p* = 3.22 × 10^−9^), malate dropped to 1/4 (Log_2_FC = –2.05, *p* = 8.30 × 10^−8^), and fumarate was similarly depleted (Log_2_FC = –1.90, *p* = 2.22 × 10^−5^) (**Figure 1, Right; Figure 2, middle row**). This simultaneous depletion of core intermediates reflects the sudden cessation of pyruvate supply from glycolysis (Ralser et al., 2007; Yun et al., 2015), demonstrating acute mitochondrial energy failure.

### 2.3. Early Signature of the Methionine Trap Coupled with Energy Depletion

We next evaluated the secondary impacts on amino acid pools. Glutamate and cysteine, key constituents for GSH synthesis and anaplerotic supply to the emptying TCA cycle, were markedly and significantly depleted in the 10 mM group (Glutamate: Log_2_FC = –0.86, *p* = 4.5 × 10^−4^; Cysteine: Log_2_FC = –0.62, *p* = 0.0027) (**Figure 2, bottom row**), reflecting rapid consumption to fuel GSH synthesis against oxidative stress (Koppula et al., 2018).

Crucially, intracellular free methionine, which was below the limit of detection (N.D.) in both control and 1 mM groups, specifically emerged and significantly accumulated 2.2-fold in the 10 mM group (Log_2_FC = 1.14, *p* = 0.011) (**Figure 2, bottom row**). Because the conversion of methionine to S-adenosylmethionine (SAM) by methionine adenosyltransferase 2A (MAT2A) strictly requires ATP (Lu, 2013), this statistically significant accumulation of methionine—occurring concurrently with acute ATP depletion and glutamate/cysteine consumption—captures the early signature of an ATP-depletion-driven “methionine trap.”

## 3. Discussion

This study revealed that the cytocidal effect of high-dose DHA on KRAS-mutant cancer cells is not merely a linear accumulation of oxidative damage, but rather an “abrupt metabolic switch” triggered when a specific concentration threshold is exceeded. Under low-dose (1 mM) DHA exposure, cells maintained overall homeostasis of glycolysis and the TCA cycle while reducing and degrading the influxed DHA. This indicates a “pre-collapse phase” where the endogenous antioxidant network possesses sufficient capacity to protect thiol-dependent enzymes such as GAPDH from ROS. Indeed, it has been reported that cells can smoothly perform reduction to ascorbic acid in response to DHA influx up to 1 mM (Abe et al., 2022).

However, at a high dose (10 mM), this DHA reduction and defense mechanism is completely saturated and breached (**Figure S1**). Our data demonstrated that within just 1 h post 10 mM exposure, the blockade of glycolysis (glycerate accumulation) and the rapid depletion of the TCA cycle (drastic decrease in α-ketoglutarate, malate and fumarate) occur simultaneously. This result strongly supports that the initial trigger of DHA-induced cell death is acute energy production failure in the mitochondria. Furthermore, the most important finding of this study is that the early signature of a “methionine trap” was confirmed in conjunction with this acute energy depletion. Under 10 mM DHA exposure, while downstream amino acids (cysteine and glutamate) are significantly depleted for GSH synthesis, the fact that methionine specifically and significantly accumulates (*p* = 0.011) demonstrates that MAT2A function is restricted by ATP depletion (Lu, 2013), physically severing the supply line of sulfur-containing amino acids that serve as “fuel” for the oxidative stress response.

## 4. Conclusion

In conclusion, the vulnerability of KRAS-mutant cells to high-dose DHA is characterized by a cascading metabolic dysfunction: the acute collapse of glycolysis and the TCA cycle, followed by a methionine trap. The elucidation of this “irreversible point of no return completed within 1 h” provides a novel biochemical foundation for the mechanisms of targeted cancer therapy using high-dose vitamin C.

## Materials and Methods

AGS cells were treated with 1 mM or 10 mM DHA for 1 h, and metabolites were extracted. Intracellular metabolomic analysis was performed using HPLC-MS/MS and GC-MS/MS. Detailed protocols for cell culture conditions, sample preparation, analytical procedures, and statistical analyses are described in the Supplementary Information.

## Acknowledgments

We would like to express our gratitude to the AI Diatrofi Research Division Subcommittee of the International Life Sciences Institute (ILSI) Japan for their financial and institutional support for this project, and to Toru Mizoguchi (Sun Chlorella Japan Corp.) for their financial assistance with the maintenance of analytical instruments. We thank Chizumi Abe (NICHe, Tohoku University) for her assistance with data collection, and Tsutomu Aida (NICHe, Tohoku University) for his administrative and logistical support. Furthermore, we are grateful to the participating corporate members of the AI Diatrofi Research Division Subcommittee, specifically Hiroshi Ochi and Hajime Nakada (Morinaga Milk Industry Co., Ltd.), Daisuke Maejima and Katsumi Kikuchi (Bourbon Corp.), Kinya Ashida (Meiji Co., Ltd.), Takahiro Matsuda (Nissin Foods Holdings Co., Ltd.), Kazuki Hattori (Yamazaki Baking Co., Ltd.), and Katsutoshi Sugimoto (ILSI Japan), for their valuable discussions and insightful feedback.

