## Supplementary Information for "Acute Metabolic Switch by High-Dose Dehydroascorbic Acid: Rapid TCA Cycle Depletion and Early Signature of Methionine Trap within 1 Hour in KRAS-Mutant Cells"

***Corresponding author:** Nozomu Hatakeyama

### Supplementary Materials and Methods

#### Reagents

Ascorbic acid (Asc), dehydroascorbic acid (DHA), and metaphosphoric acid (MPA) were purchased from FUJIFILM Wako Pure Chemical Corporation (Osaka, Japan). Fetal bovine serum (FBS), RPMI 1640 medium (Cat. R8758-500ML), and penicillin-streptomycin solution (Cat. P4458-100ML) were purchased from Sigma-Aldrich (St. Louis, MO, USA). Trypan blue solution (0.4%; Cat. 15250061) and a bicinchoninic acid (BCA) protein assay kit were purchased from Thermo Fisher Scientific Inc. (Waltham, MA, USA). All other chemicals were of analytical reagent grade and were used without further purification.

#### Routine Cell Culture

Human gastric adenocarcinoma cells (AGS cells) were obtained from the American Type Culture Collection (ATCC) and maintained in RPMI 1640 medium supplemented with 10% (v/v) FBS and 1% (v/v) penicillin–streptomycin solution (hereafter referred to as complete medium). For routine culture, cells were seeded at a density of $1.0 \times 10^5$ cells/mL in 10-cm culture dishes.

**DHA Treatment and Quantification of Intracellular Ascorbic Acid**

AGS cells were seeded in 6-well plates at a density of $2.0 \times 10^4$ cells/well and cultured until sub-confluent, with medium replacement every 2 days. Cells were then incubated in complete medium containing 0, 0.2, 1, or 10 mM DHA for 60 min. At each time point, intracellular Asc was extracted using a procedure based on the method described by Abe et al. [1]. Intracellular Asc was quantified by high-performance liquid chromatography coupled with tandem mass spectrometry (HPLC-MS/MS; QTRAP 6500; SCIEX, Framingham, MA, USA) [1]. Cell counts were determined using trypan blue staining.

**DHA Treatment and Analysis of Intracellular Metabolites**

AGS cells were seeded in 6-well plates at a density of $2.0 \times 10^4$ cells/well and cultured until sub-confluent. Cells were then incubated in complete medium containing 1 mM or 10 mM DHA for 60 min. At the designated time point, intracellular metabolites were extracted and derivatized according to the protocols provided with the Smart Metabolites Database (Shimadzu Corporation, Kyoto, Japan) [2]. Samples were analyzed by gas chromatography-tandem mass spectrometry (GC-MS/MS; GCMS-TQ8040; Shimadzu Corporation), as described previously. To minimize compositional changes, sample extraction, derivatization, and instrumental analysis were performed sequentially without intermediate storage.

For each metabolite, the relative abundance was calculated as the peak area ratio of the target metabolite to the internal standard (2-isopropylmalic acid). Cellular protein concentrations were determined using a BCA protein assay to normalize the metabolite levels.

**Statistical Analysis**

Statistical analyses were performed using Python (SciPy library). For metabolomics data, the relative intracellular levels between the control group and DHA-treated groups (1 mM, $n = 11$ from 2 independent batches; 10 mM, $n = 18$ from 3 independent batches) were compared using a two-sided Welch's $t$-test on $\log_2$-transformed values, which does not assume equal variances. For other biochemical assays (e.g., Figure S1), statistical differences were evaluated using a two-sided Welch's t-test on raw relative values. A $p$-value of < 0.05 was considered statistically significant.

### Supplementary Figures

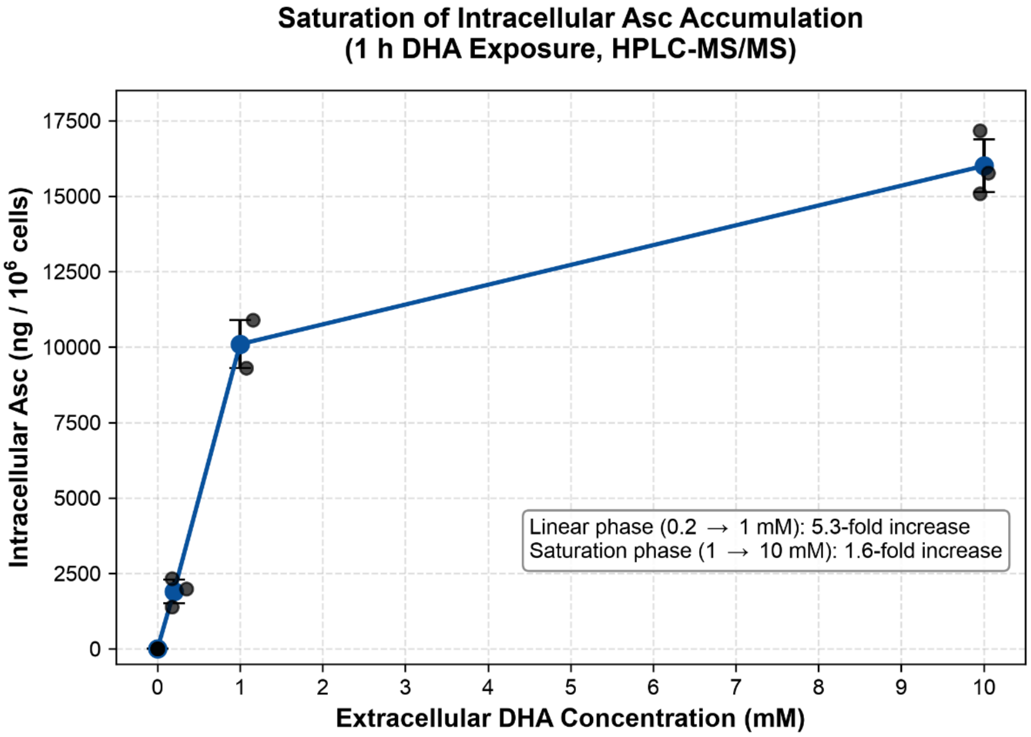


**Figure S1. Saturation of intracellular ascorbic acid (Asc) accumulation under high-dose DHA exposure in AGS cells.**

AGS cells were exposed to 0, 0.2, 1, or 10 mM DHA for 60 min, and intracellular Asc concentrations were quantified using HPLC-MS/MS [1]. Data represent mean ± SD ($n = 3$ for 0, 0.2, and 10 mM groups; $n = 2$ for 1 mM group, excluding one technical extraction failure). While increasing extracellular DHA from 0.2 mM to 1 mM led to a linear 5.31-fold increase in intracellular Asc (1,900.0 to 10,092.9 ng/$10^6$ cells), which is consistent with the linear uptake dynamics at lower concentrations (< 1 mM) previously reported [1], a 10-fold increase in extracellular DHA from 1 mM to 10 mM resulted in only a 1.59-fold increase (16,000.0 ng/$10^6$ cells; Welch's $t$-test, $p = 0.023$). This saturation curve indicates that the intracellular reductive capacity (e.g., GSH-dependent reduction) is overwhelmed at 10 mM, triggering the accumulation of unreduced DHA and its subsequent non-enzymatic degradation.

[2] Shimadzu Corporation, Smart Metabolites Database: database software for GC/MS and GC-MS/MS analysis of metabolites, Shimadzu Corporation, Kyoto, Japan, C146-E277A, 2018.
